# Chemoproteomic profiling of 8-oxoguanosine-sensitive RNA-protein interactions

**DOI:** 10.1101/2023.09.01.555928

**Authors:** Jennifer Villers, Eliana McCann Smith, Amanda N. DeLiberto, A. Emilia Arguello, Joy Nyaanga, Ralph E. Kleiner

## Abstract

Cellular nucleic acids are subject to assault by endogenous and exogenous agents that can perturb the flow of genetic information. Oxidative stress leads to the accumulation of 8-oxoguanine (8OG) on DNA and RNA. 8OG lesions on mRNA negatively impact translation, but their effect on global RNA-protein interactions is largely unknown. Here, we apply an RNA chemical proteomics approach to investigate the effect of 8OG on RNA-protein binding. We find proteins that bind preferentially to 8OG-modified RNA, including IGF2BP1-3 and hnRNPD, and proteins that are repelled by 8OG such as RBM4. We characterize these interactions using biochemical and biophysical assays to quantify the effect of 8OG on binding and show that a single 8OG abolishes binding of RBM4 to its preferred CGG-containing substrate. Taken together, our work establishes the molecular consequences of 8OG on cellular RNA-protein binding and provides a framework for interrogating the role of RNA oxidation in biological systems.

## INTRODUCTION

Nucleic acids are continuously exposed to damaging agents, such as reactive oxygen species (ROS) generated through normal cellular metabolism, that can alter their chemical structure and function. The reaction of ROS with DNA and RNA can result in numerous modifications, the most prevalent of which is the oxidation of guanine to generate 7,8-dihydro-8-oxoguanine (8OG or 8-oxoG)^1, 2^. 8OG in DNA is a well-studied mutagenic lesion that is recognized and processed by DNA repair pathways (reviewed in ^3, 4^). In contrast, relatively little is known about the biological consequences of RNA oxidation and the interaction of oxidative lesions such as 8OG with RNA-associated processes.

RNA oxidation is widespread in cells and associated with pathophysiological processes (reviewed in ^5-8^). Brain tissue analyzed from patients with neurodegenerative diseases (including Alzheimer’s, Parkinson’s, and amyotrophic lateral sclerosis) contains higher levels of 8OG in cytoplasmic and nuclear RNA as compared to healthy individuals^9-14^. Interestingly, 8OG levels are higher in RNA than in DNA, perhaps because RNA is more single-stranded and more weakly associated with proteins, which exposes the nucleobases to attack by ROS^10, 15-17^. Although it remains unclear whether the accumulation of 8OG in RNA is a precursor or a consequence of neurodegeneration, the inability to effectively clear damaged RNA may contribute to disease pathogenicity^18^.

Functional studies of 8OG lesions in RNA have primarily focused on its deleterious translational effects. *In vitro* oxidized mRNA transfected into mammalian cells was found to produce less functional protein than the corresponding nonoxidized mRNA^19^, consistent with errors in translational efficiency and fidelity. Similar findings were reported in cells fed 8-oxo-GTP^20^. Using a reconstituted bacterial translation system, Zaher and co-workers showed that a single 8OG lesion can decrease the rate of peptide-bond formation for cognate and near-cognate aminoacyl-tRNA by >1000-fold^21^. These data highlight the detrimental effect of 8OG in mRNA on translation and suggest that the ribosome may play an important role in the surveillance and clearance of oxidized transcripts through the No-Go Decay (NGD) pathway^21, 22^.

In addition to interactions with the ribosome, mRNAs engage with a host of RNA-binding proteins (RBPs) that regulate all aspects of RNA behavior^23^ and whose binding may be sensitive to the presence of 8OG. A number of RBPs have been reported to bind 8OG-containing RNA including Y-box binding protein 1 (YBX1)^24^, human polynucleotide phosphorylase protein (hPNPase)^25^, heterogenous nuclear ribonucleoprotein D (hnRNPD/AUF1)^26^, and poly(C)-binding protein (PCBP1)^27^. Specific recognition of 8OG-RNA by these proteins has been proposed to play a role in the metabolism of oxidized RNA as well as cell fate decisions, however most were identified using heavily oxidized RNAs containing multiple 8OG modifications, which is unlikely to reflect the physiological abundance of 8OG in the transcriptome (estimated at 1 in 10^5^ nucleotides in H_2_O_2_-treated cells^16^). Further, quantitative biophysical analysis of the binding preference of these proteins for a single 8OG lesion within an RNA strand is lacking, and there is little known about the ability of 8OG lesions to abrogate RBP-RNA recognition events.

Here, we apply an RNA chemical proteomics approach previously developed in our group^28, 29^ to investigate the binding of cellular proteins to oligonucleotides containing single 8OG lesions. We identify several RBPs that preferentially recognize 8OG-modified RNA including hnRNPD and proteins from the IGF2BP family. In addition, we find RBPs that are strongly repelled by 8OG such as RNA binding motif protein 4 (RBM4). We characterize the identified RNA-protein interactions using *in vitro* biochemical and biophysical assays to quantify the effect of 8OG and establish dependence on surrounding RNA sequence context. Taken together, our work provides a molecular framework for studying RNA oxidation and its effect on RNA-associated regulatory proteins.

## RESULTS

To identify the effect of 8OG RNA lesions on global RNA-protein interactions, we applied a comparative RNA chemical proteomics method developed by our group that relies on diazirine-modified oligonucleotide probes and quantitative mass spectrometry^28, 29^. In contrast to enzyme-dependent RNA modifications that typically occur within a preferred consensus sequence, 8OG lesions are thought to be produced by non-enzymatic oxidation processes and likely exhibit only minimal sequence bias as determined by oxidation potential. Therefore, we designed synthetic 10 nt RNA probes with arbitrary sequence containing (Fig. 1A): (i) a single 8OG or G residue at a defined position; (ii) a photocrosslinkable diazirine-modified uridine residue (5-DzU); and (iii) 3’-biotin modification (Table 1, Supplementary Table 1). We photocrosslinked HeLa cell lysates with 8OG (**1**) or G (**2**) oligo probes. Crosslinked protein-RNA complexes were then isolated by streptavidin enrichment, eluted from the beads with mild RNase treatment, and analyzed by label-free LC-MS/MS to quantify and compare the abundance of proteins crosslinked to 8OG or G probes. Three biological replicates were performed and results are displayed in a volcano plot (Fig. 1B, Supplementary Dataset 1).

**Table 1.** Oligonucleotides used in this work.

| Oligo | RNA Sequence |
| --- | --- |
| 1 | 5'-AC-8OG-UG-5DzU-GUAC-biotin-3' |
| 2 | 5'-ACGUG-5DzU-GUAC-biotin-3' |
| 3 | 5'-GA-8OG-CU-5DzU-GUAC-biotin-3' |
| 4 | 5'-GAGCU-5DzU-GUAC-biotin-3' |
| 5 | 5'-AC-8OG-UGUGUAC-fluorescein-3' |
| 6 | 5'-ACGUGUGUAC-fluorescein-3' |
| 7 | 5'-GUAAC-8OG-GUAC-fluorescein-3' |
| 8 | 5'-GUAACGGUAC-fluorescein-3' |
| 9 | 5'-CGCG-8OG-CGGUA-fluorescein-3' |
| 10 | 5'-CGCGGCGGUA-fluorescein-3' |

**Figure 1.**
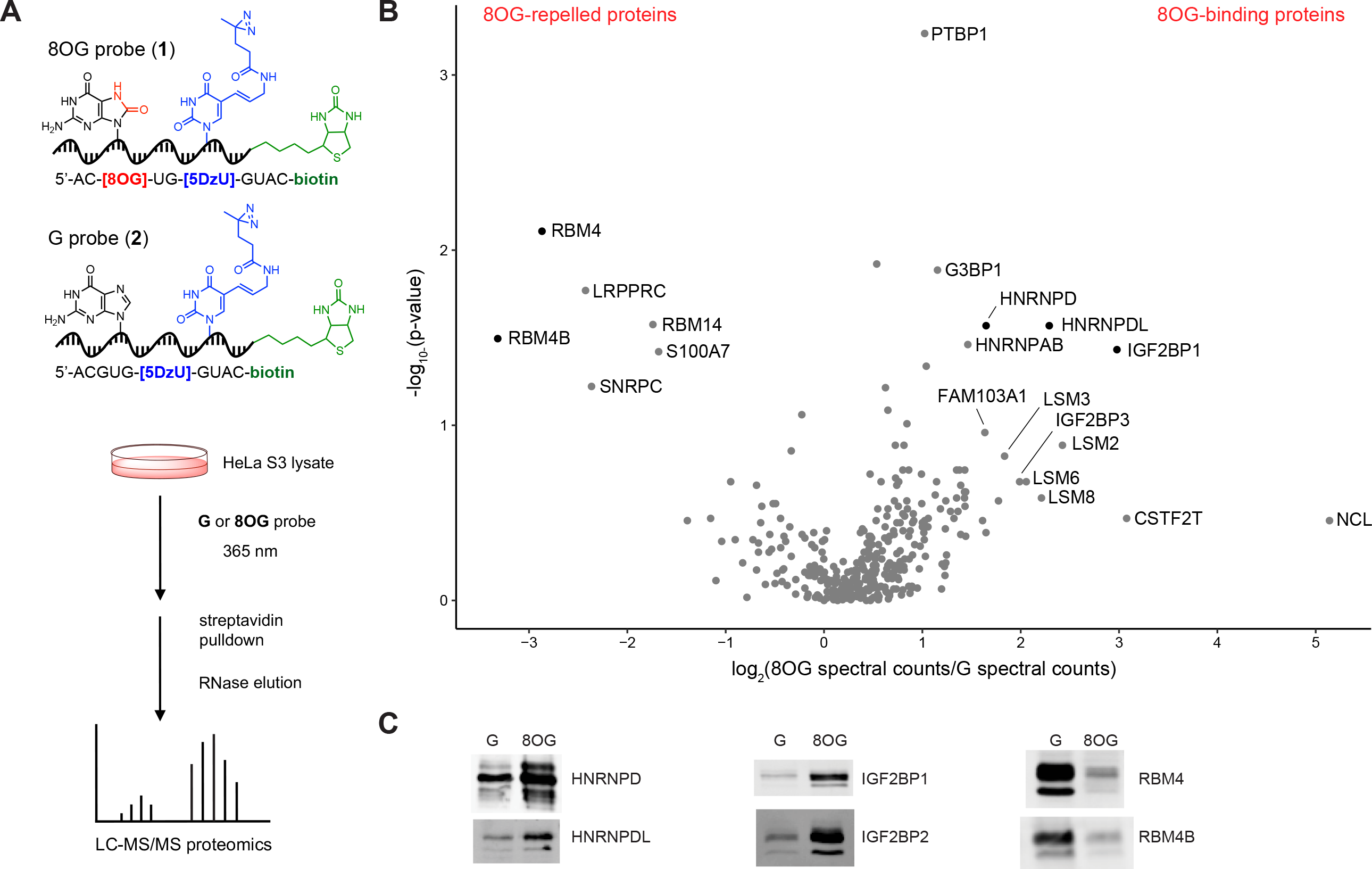
Proteomic profiling of 8OG-RNA interactome. **(A)** Structure of 8OG or G oligo probes. Schematic for chemical proteomics workflow. **(B)** Volcano plot of protein enrichment ratios (8OG/G spectral counts) and p-values from label-free proteomics experiments (n=3). To calculate enrichment ratios for proteins identified in only one of the two conditions (G or 8OG probe), we added 1 to all spectral count values. P-values were calculated based on a student’s t-test. **(C)** Validation of protein hits from **(B)**. Lysates were photocrosslinked with 8OG (**1**) or G (**2**) oligo probe and RBPs were detected by western blotting after streptavidin enrichment. IGF2BP1 and hnRNPD were detected with corresponding antibodies against endogenous protein. Anti-FLAG M2 antibody was used to detect overexpressed epitope-tagged hnRNPDL, IGF2BP2, RBM4, and RBM4B. See Supplementary Fig. 1 for full western blotting data.

We identified ∼340 proteins of which most showed minimal preference for crosslinking to either 8OG or G probe. Proteins exhibiting fold-change greater than 2.5 and p-value<0.05 were considered selective G or 8OG binders and investigated further. Using these criteria, we identified four potential 8OG-binding proteins. Among them, the heterogenous nuclear ribonucleoproteins hnRNPAB, hnRNPD and hnRNPDL showed between 2.7-fold to 5-fold preference for the 8OG probe (Fig. 1B). HnRNP proteins are a large family of RNA-binding proteins involved in mRNA trafficking and stability, alternative splicing, and transcriptional and translational regulation^30^. Gratifyingly, hnRNPD/AUF1 has been previously identified as an 8OG binder^26, 31^ indicating that our method can reliably capture known 8OG-interacting RBPs. In addition, we found insulin-like growth factor 2 mRNA-binding protein 1 (IGF2BP1), which demonstrated ∼8-fold preference for the 8OG probe. The related proteins IGFBP2 and IGFBP3 were also enriched in our dataset, although they did not meet our criteria for significance (Supplementary Dataset 1). IGF2BP proteins are a conserved family of RNA-binding proteins with roles in mRNA processing, localization, translation, and stability^32^. We did not observe enrichment of PCBP1 or PCBP2 in our dataset with the 8OG probe. PCBP1 has been reported to bind heavily oxidized RNA^27^, but not oligonucleotides containing a single 8OG lesion, such as our probe. YBX1 and hPNPase (PNPT1) have also been reported as 8OG-binding proteins^24, 25^; we did not observe significant enrichment of YBX1 in our data and PNPT1 was not identified. In addition to putative 8OG binding proteins, we also identified proteins enriched by the G probe (Fig. 1B). Among them, the RNA-binding proteins RBM4 and RBM4B, which are involved in alternative splicing and translational regulation^33^, showed between 7-fold to 10-fold preference for the G probe. We confirmed our proteomics hits using photocrosslinking, pulldown, and western blotting (Fig. 1C, Supplementary Fig. 1), which showed the expected trends – hnRNPD, hnRNPDL, IGF2BP1 and IGF2BP2 preferentially bind to the 8OG probe while RBM4 and RBM4B have a strong preference for the G probe. Additionally, to interrogate the sequence specificity of 8OG recognition by IGF2BP1 and hnRPND, we synthesized diazirine-containing oligo probes **3** and **4** with scrambled sequence (Table 1, Supplementary Table 1) and assayed binding by photocrosslinking, pulldown, and western blotting. IGF2BP1 and HNRPND crosslinked preferentially to the scrambled 8OG probe (**3**) over the scrambled G probe (**4**), indicating that recognition of an 8OG lesion by these proteins is not sequence-specific (Supplementary Fig. 2).

To biochemically characterize the interaction of IGF2BP1 with 8OG-modified RNA, we generated recombinant proteins and measured *in vitro* binding or crosslinking. IGF2BP proteins contain six RNA-binding domains: two RNA-recognition motifs (RRM) and four K homology (KH) domains. The KH domains have been shown to be the major RNA-binding modules^34^. Since full-length human IGF2BP1 is difficult to purify from *E. coli*, we separately purified proteins comprising the N-terminal portion of IGF2BP1 containing two RRMs (amino acids 1-194, hereafter termed “IGF2BP1-RRM”), and the C-terminal portion containing four KH domains (amino acids 195-577, hereafter termed “IGF2BP1-KH”) (Fig. 2A, Supplementary Fig. 3A). We determined the photocrosslinking efficiency for IGF2BP1-KH and IGF2BP1-RRM by performing dose titrations with oligo probe pairs **1**/**2** and **3**/**4**. We found that crosslinking to IGF2BP1-RRM was less efficient than to IGF2BP1-KH (Supplementary Fig. 3), consistent with weak RRM-RNA binding demonstrated by prior work on IGF2BP homologs^34^, and therefore we primarily focused on IGF2BP1-KH. Photocrosslinking of IGF2BP1 with oligo probe pairs **1**/**2** or **3**/**4** demonstrated clear preference for the 8OG-containing probes (**1** and **3**) with ∼4-fold to ∼8-fold lower EC_50_ (Supplementary Fig. 3B, 3C). As crosslinking is not an equilibrium measurement, we next measured binding affinity of IGF2BP1-KH for unmodified RNA or 8OG-containing RNA by fluorescence polarization (FP) and electrophoretic mobility shift assay (EMSA) using fluorescein-labeled oligonucleotides (**5**/**6**) with the same sequence as our original photocrosslinkable probes **1**/**2** (Table 1, Supplementary Table 1). Both equilibrium binding assays demonstrated a 1.68-fold to 2-fold binding preference of IGF2BP1-KH for 8OG-modified oligo **5** (Fig. 2A, Fig. 2B, Supplementary Fig. 4), with K_d_ values of 275 ± 13 nM (FP) and 304 ± 53 nM (EMSA) for oligo **5** and 462 ± 11 nM (FP) and 635 ± 113 nM (EMSA) for the G oligo (**6**).

**Figure 2.**
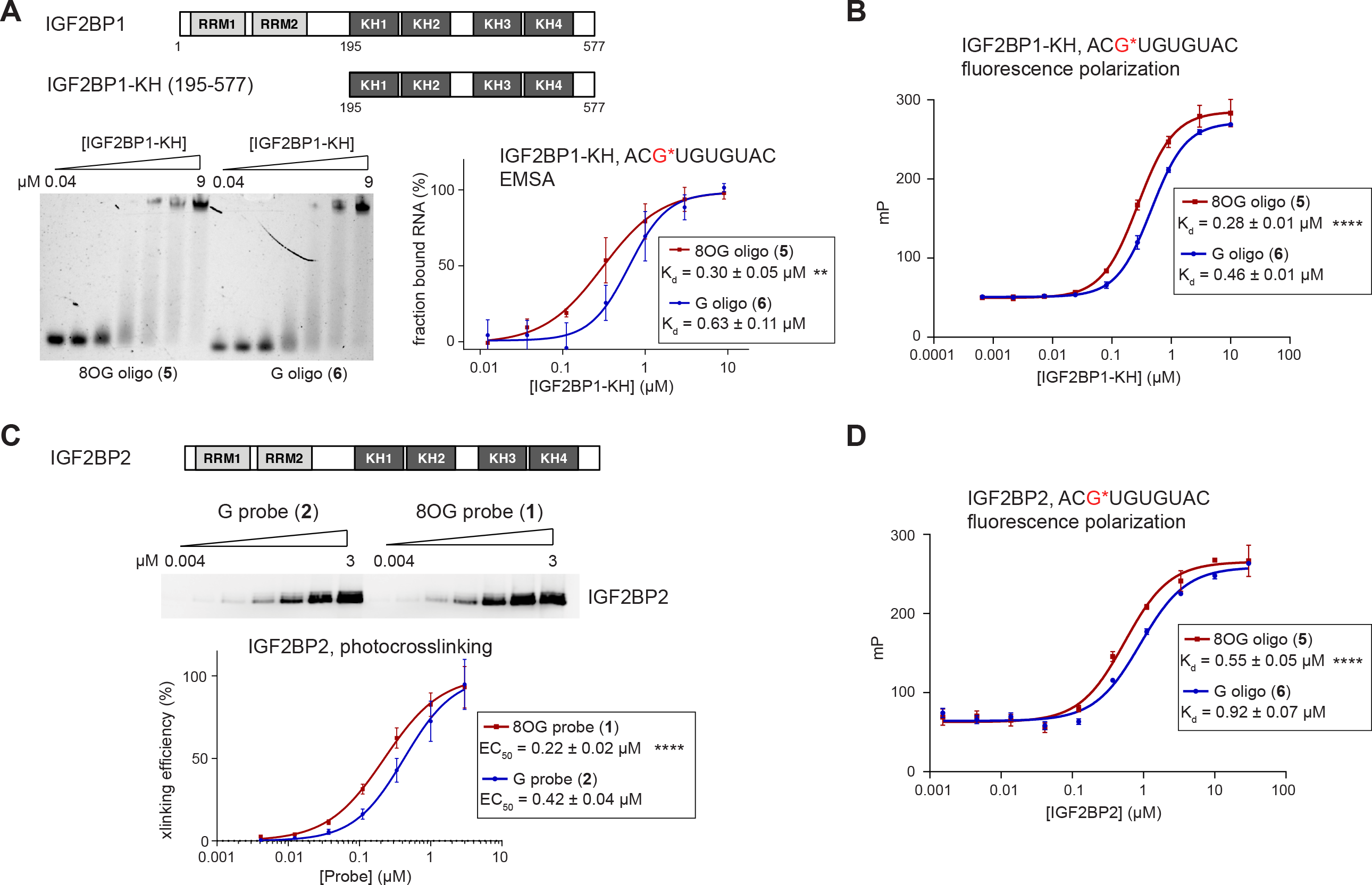
Biochemical characterization of interaction between IGF2BP1/2 and 8OG-containing RNA oligos. **(A)** Electrophoretic mobility shift assay (EMSA) with IGF2BP1-KH (195-577) and fluorescein-labeled 8OG (**5**) or G (**6**) oligos. Oligos were mixed with various concentrations of IGF2BP1 and RNA-protein complexes were resolved on a TGE native gel and detected using in-gel fluorescence. See Supplementary Fig. 3 for full EMSA data. K_d_ values were determined by fitting data to a sigmoidal dose-response curve. Values represent mean ± s.e. (n=3). Asterisks represent statistically significant differences in binding affinity (**, p<0.01). **(B)** Fluorescence anisotropy assay with IGF2BP1-KH and oligos **5** and **6**. Oligos were mixed with various concentrations of protein and binding was monitored by fluorescence polarization. K_d_ values were determined by fitting data to a sigmoidal dose-response curve. Values represent mean ± s.e. (n=4). Asterisks represent statistically significant differences in binding affinity (****, p<0.0001). **(C)** Photocrosslinking of IGF2BP2 with 8OG (**1**) or G (**2**) diazirine-containing oligo probes. IGF2BP2 was mixed with various concentrations of probe and UV irradiated at 365 nm. Crosslinked RNA-protein complexes were detected by streptavidin western blotting. See Supplementary Fig. 4 for full blot. EC_50_ values were determined by fitting data to a sigmoidal dose-response curve. Values represent mean ± s.e. (n=4). Asterisks represent statistically significant differences in crosslinking (****, p<0.0001). **(D)** Fluorescence polarization assay with IGF2BP2 and 8OG (**5**) or G (**6**) oligos. Experiment was performed as described in **(B)**. Values represent mean ± s.e. (oligo **5**, n=4; oligo **6**, n=3;).

In the absence of full-length IGF2BP1 and without the certitude that the KH domains fully recapitulate the binding affinity of the full-length protein, we decided to express and purify full-length IGF2BP2 and characterize its binding to 8OG/G oligos, since IGF2BP1 and IGF2BP2 share high sequence and structural homology. IGF2BP2 binding affinity was first characterized by photocrosslinking titrations with oligo probes **1**/**2**. We measured a 1.9-fold difference in EC_50_ between the two reactions: 224 ± 15 nM with 8OG probe **1** and 424 ± 36 nM with G probe **2** (Fig. 2C, Supplementary Fig. 5). Further, we assayed photocrosslinking with the scrambled probes **3**/**4** and found a ∼7-fold preference for the 8OG probe (**3**) over the corresponding G probe (**4**) (Supplementary Fig. 6). We confirmed the preference of full-length IGF2BP2 for 8OG-modified RNA by fluorescence polarization binding assay with oligos **5**/**6** measuring a K_d_ of 546 ± 47 nM with 8OG oligo **5** and 920 ± 70 nM with G oligo **6** (Fig. 2D). The preference of IGF2BP proteins for binding to 8OG-modified RNA is consistent across multiple assays, protein family members and constructs, and RNA sequences indicating that these proteins can be considered putative 8OG reader proteins.

Our proteomics experiment also identified proteins that are repelled by 8OG and we therefore explored the binding properties of the top hit, RBM4 (Fig. 1B, 1C). Since we had difficulties generating full-length RBM4 through heterologous expression in *E. coli*, we focused on studying a truncated construct containing the two RRM domains and CCHC Zinc finger motif of RBM4 (amino acids 1-176), which should primarily mediate RNA binding. We first analyzed binding affinity by photocrosslinking titrations with oligo probes **1**/ **2** which demonstrated >10-fold preference for the G probe **2** over 8OG probe **1** (Fig. 3A, Supplementary Fig. 7), consistent with our proteomics and western blot data. Further, we confirmed the strong repulsive effect of 8OG on RBM4 binding using fluorescence polarization and observed no detectable binding to the 8OG oligo (**5**), whereas we measured a K_d_ of 620 ± 41 nM for the G oligo (**6**) (Fig. 3B). Native RNA binding sites of RBM4 are enriched in CGG motifs^35-38^, therefore we introduced an 8OG lesion into two different CGG-containing oligonucleotide sequences (Table 1, Supplementary Table 1) and assayed RBM4 binding by fluorescence polarization. Consistent with our previous observations, RBM4 did not bind to oligo **7**, which contains the motif C(8OG)G, whereas it bound to the corresponding unoxidized oligo (**8**) with K_d_ = 5.28 ± 1.27 µM (Fig. 3C). We observed overall stronger binding to a set of oligos containing tandem CGG motifs (i.e. CGG*CGG) (oligos **9**/**10**) with a K_d_ of 56 ± 5 nM for the unoxidized oligo (**10**) and K_d_ of 86 ± 12 nM for the 8OG oligo (**9**) (Fig. 3D). In this sequence context, 8OG only slightly perturbed RBM4 binding, likely due to the presence of a single 8OG lesion, leaving one CGG motif unoxidized. Overall, using three different sequence contexts, we consistently show that RBM4-RNA interactions are inhibited by a single 8OG lesion.

**Figure 3.**
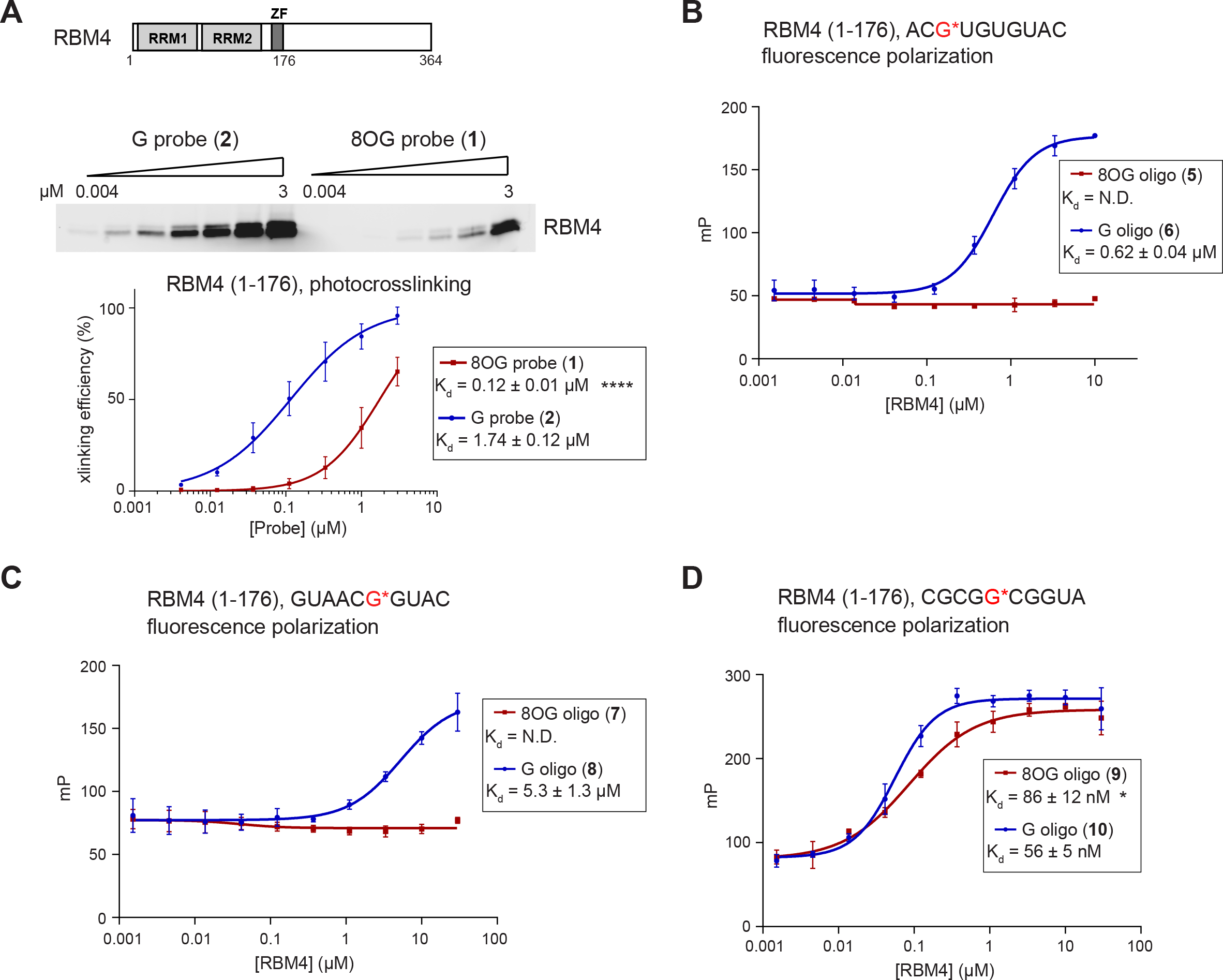
Biochemical characterization of interaction between RBM4 and 8OG-containing RNA oligos. **(A)** Photocrosslinking of RBM4-RRM (1-176) with the 8OG (**1**) or G (**2**) oligo probes. Experiment was performed as in Fig. 2C. See Supplementary Fig. 6 for full blot. EC_50_ values were determined by fitting data to a sigmoidal dose-response curve. Values represent mean ± s.e. (n=4). Asterisks represent statistically significant differences in binding affinity (****, p<0.0001). **(B)** Fluorescence polarization assay with RBM4-RRM and oligos **5** and **6**. Experiment was performed as in Fig. 2B. Values represent mean ± s.e. (oligo **5**, n=3; oligo **6**, n=5). **(C)** Fluorescence polarization assay with RBM4-RRM and oligos **7** and **8** containing the following nucleotide sequence: 5’-GUAACG*GUAC (G* = 8OG or G). Experiment was performed as described in Fig. 2B. Values represent mean ± s.e. (oligo **7**, n=2; oligo **8**, n=4). **(D)** Fluorescence polarization assay with RBM4-RRM and oligos **9** and **10** containing the following nucleotide sequence: CGCGG*CGGUA (G* = 8OG or G). Experiment was performed as described in Fig. 2B. Values represent mean ± s.e. (n=4). Asterisk represents statistically significant differences in binding affinity (*, p<0.05).

## DISCUSSION

In this manuscript, we profile the effects of 8OG on cellular RNA-protein interactions. Whereas most RBP-RNA binding events detected in our chemoproteomics study are unaffected by the presence of a single 8OG lesion, we find a small set of RBPs showing 8OG-sensitive binding behavior. In particular, we show that IGF2BP family proteins bind more tightly to 8OG-modified RNA across a range of sequence contexts, implicating these proteins as 8OG ‘readers’. We also identify hnRNPD, which was previously shown to interact with oxidized RNA^26^, as an 8OG-binding protein. In contrast, a single 8OG lesion can abrogate RBM4 binding in its native CGG recognition motif. Previous work has identified PCBP1^27^, hPNPase^25^, and YBX1^24^ as 8OG-RNA binding proteins, but we do not observe significant enrichment of these proteins with our 8OG oligonucleotide probe, possibly due to sequence bias or only modest preference for a single 8OG residue. Our work expands our understanding of the effect of RNA oxidation on global RNA-protein interactions.

The binding of hnRNPD to 8OG-RNA has been proposed as a quality control mechanism to selectively degrade oxidized RNA^39^. This function is analogous to the role of hnRNPD in destabilizing mRNAs with AU-rich elements^40^. IGF2BP proteins are known to regulate RNA stability, translation, and localization^32^, however, in contrast to hnRNPD, IGF2BP binding is associated with mRNA stability. These seemingly disparate functions are challenging to reconcile within the context of RNA oxidation. Interestingly, IGF2BP1/2 and hnRNPD have been shown to interact with one another^41, 42^. Further investigation of the biology of these proteins in the context of 8OG-modified RNA will be necessary to decipher their relevance to RNA oxidation.

Although most studies of 8OG on RNA have focused on the identification of proteins that bind this lesion, our work identifies the opposite behavior – i.e. an RBP that cannot bind to its oxidized substrate sequence. We show that RBM4 is uniquely sensitive to 8OG lesions within multiple sequence contexts including its preferred CGG binding motif. RBM4 has diverse roles in RNA biology including alternative splicing^43-45^, translation regulation^35, 46^, and RNA stability^38^ and evidence suggests that its activity is regulated by oxidative stress. Our biochemical study supports the idea that these processes would be perturbed upon guanine oxidation occurring within the RBM4 recognition site. Whereas this phenomenon may reflect the ability of RNA oxidation to perturb normal post-transcriptional regulation, an alternative hypothesis is that RBM4 sites, which are typically G-rich and therefore more prone to 8OG formation, may have evolved to function as sensors of oxidative stress. Similarly, we propose that RBPs with G-rich binding sites should be particularly sensitive to the effects of 8OG in the transcriptome. Our study primarily identified RBM4/4B, but other 8OG-sensitive interactions may be revealed by altering the sequence of the oligonucleotide probe. Further, future work should focus on identifying endogenous 8OG sites that are upregulated upon ROS production and connecting these lesions to functional RNA-protein interactions to understand the effect of oxidation on native RNA regulation.

## ACKNOWLEDGMENTS

We are grateful to Joan Steitz for the kind gift of anti-HNRNPD antibody and Stefan Huttelmaier for the kind gift of anti-IGF2BP1 antibody and for advice on IGF2BP1/2 protein expression and purification. We thank Tharan Sirakumar, Saw Kyin, and Henry Shwe at the Princeton University Mass Spectrometry Core Facility for performing proteomics analysis. We thank Hahn Kim and the Princeton University Small Molecule Screening Facility for assistance with fluorescence polarization assays. R.E.K. acknowledges support from the National Institutes of Health (R01 GM132189). A.E.A was supported by an Edward C. Taylor 3rd Year Graduate Fellowship in Chemistry. All authors acknowledge financial support from Princeton University.

## REFERENCES

(1) Cadet, J.; Wagner, J. R. DNA base damage by reactive oxygen species, oxidizing agents, and UV radiation. Cold Spring Harb Perspect Biol 2013, 5 (2). DOI: 10.1101/cshperspect.a012559 From NLM Medline.

(2) Kong, Q.; Lin, C. L. Oxidative damage to RNA: mechanisms, consequences, and diseases. Cell Mol Life Sci 2010, 67 (11), 1817–1829. DOI: 10.1007/s00018-010-0277-y From NLM Medline.

(3) Lindahl, T.; Wood, R. D. Quality control by DNA repair. Science 1999, 286 (5446), 1897–1905.

(4) Barnes, D. E.; Lindahl, T. Repair and genetic consequences of endogenous DNA base damage in mammalian cells. Annu Rev Genet 2004, 38, 445–476. DOI: 10.1146/annurev.genet.38.072902.092448.

(5) Li, Z.; Wu, J.; Deleo, C. J. RNA damage and surveillance under oxidative stress. IUBMB Life 2006, 58 (10), 581–588. DOI: 10.1080/15216540600946456.

(6) Wurtmann, E. J.; Wolin, S. L. RNA under attack: cellular handling of RNA damage. Crit Rev Biochem Mol Biol 2009, 44 (1), 34–49. DOI: 10.1080/10409230802594043.

(7) Li, Z.; Malla, S.; Shin, B.; Li, J. M. Battle against RNA oxidation: molecular mechanisms for reducing oxidized RNA to protect cells. Wiley Interdiscip Rev RNA 2014, 5 (3), 335–346. DOI: 10.1002/wrna.1214.

(8) Simms, C. L.; Zaher, H. S. Quality control of chemically damaged RNA. Cell Mol Life Sci 2016, 73 (19), 3639–3653. DOI: 10.1007/s00018-016-2261-7.

(9) Zhang, J.; Perry, G.; Smith, M. A.; Robertson, D.; Olson, S. J.; Graham, D. G.; Montine, T. J. Parkinson’s disease is associated with oxidative damage to cytoplasmic DNA and RNA in substantia nigra neurons. Am J Pathol 1999, 154 (5), 1423–1429. DOI: 10.1016/S0002-9440(10)65396-5.

(10) Nunomura, A.; Perry, G.; Pappolla, M. A.; Wade, R.; Hirai, K.; Chiba, S.; Smith, M. A. RNA oxidation is a prominent feature of vulnerable neurons in Alzheimer’s disease. J Neurosci 1999, 19 (6), 1959–1964.

(11) Chang, Y.; Kong, Q.; Shan, X.; Tian, G.; Ilieva, H.; Cleveland, D. W.; Rothstein, J. D.; Borchelt, D. R.; Wong, P. C.; Lin, C. L. Messenger RNA oxidation occurs early in disease pathogenesis and promotes motor neuron degeneration in ALS. PLoS One 2008, 3 (8), e2849. DOI: 10.1371/journal.pone.0002849.

(12) Nunomura, A.; Chiba, S.; Kosaka, K.; Takeda, A.; Castellani, R. J.; Smith, M. A.; Perry, G. Neuronal RNA oxidation is a prominent feature of dementia with Lewy bodies. Neuroreport 2002, 13 (16), 2035–2039.

(13) Shan, X.; Tashiro, H.; Lin, C. L. The identification and characterization of oxidized RNAs in Alzheimer’s disease. J Neurosci 2003, 23 (12), 4913–4921.

(14) Bradley-Whitman, M. A.; Timmons, M. D.; Beckett, T. L.; Murphy, M. P.; Lynn, B. C.; Lovell, M. A. Nucleic acid oxidation: an early feature of Alzheimer’s disease. J Neurochem 2014, 128 (2), 294–304. DOI: 10.1111/jnc.12444.

(15) Fiala, E. S.; Conaway, C. C.; Mathis, J. E. Oxidative DNA and RNA damage in the livers of Sprague-Dawley rats treated with the hepatocarcinogen 2-nitropropane. Cancer Res 1989, 49 (20), 5518–5522.

(16) Hofer, T.; Badouard, C.; Bajak, E.; Ravanat, J. L.; Mattsson, A.; Cotgreave, I. A. Hydrogen peroxide causes greater oxidation in cellular RNA than in DNA. Biol Chem 2005, 386 (4), 333–337. DOI: 10.1515/BC.2005.040 From NLM Medline.

(17) Hofer, T.; Seo, A. Y.; Prudencio, M.; Leeuwenburgh, C. A method to determine RNA and DNA oxidation simultaneously by HPLC-ECD: greater RNA than DNA oxidation in rat liver after doxorubicin administration. Biol Chem 2006, 387 (1), 103–111. DOI: 10.1515/BC.2006.014.

(18) Shan, X.; Chang, Y.; Lin, C. L. Messenger RNA oxidation is an early event preceding cell death and causes reduced protein expression. FASEB J 2007, 21 (11), 2753–2764. DOI: 10.1096/fj.07-8200com.

(19) Tanaka, M.; Chock, P. B.; Stadtman, E. R. Oxidized messenger RNA induces translation errors. Proc Natl Acad Sci U S A 2007, 104 (1), 66–71. DOI: 10.1073/pnas.0609737104 From NLM Medline.

(20) Dai, D. P.; Gan, W.; Hayakawa, H.; Zhu, J. L.; Zhang, X. Q.; Hu, G. X.; Xu, T.; Jiang, Z. L.; Zhang, L. Q.; Hu, X. D.; Nie, B.; Zhou, Y.; Li, J.; Zhou, X. Y.; Li, J.; Zhang, T. M.; He, Q.; Liu, D. G.; Chen, H. B.; Yang, N.; Zuo, P. P.; Zhang, Z. X.; Yang, H. M.; Wang, Y.; Wilson, S. H.; Zeng, Y. X.; Wang, J. Y.; Sekiguchi, M.; Cai, J. P. Transcriptional mutagenesis mediated by 8-oxoG induces translational errors in mammalian cells. Proc Natl Acad Sci U S A 2018, 115 (16), 4218–4222. DOI: 10.1073/pnas.1718363115 From NLM Medline.

(21) Simms, C. L.; Hudson, B. H.; Mosior, J. W.; Rangwala, A. S.; Zaher, H. S. An active role for the ribosome in determining the fate of oxidized mRNA. Cell Rep 2014, 9 (4), 1256–1264. DOI: 10.1016/j.celrep.2014.10.042.

(22) Doma, M. K.; Parker, R. Endonucleolytic cleavage of eukaryotic mRNAs with stalls in translation elongation. Nature 2006, 440 (7083), 561–564. DOI: 10.1038/nature04530.

(23) Gerstberger, S.; Hafner, M.; Tuschl, T. A census of human RNA-binding proteins. Nat Rev Genet 2014, 15 (12), 829–845. DOI: 10.1038/nrg3813 From NLM Medline.

(24) Hayakawa, H.; Uchiumi, T.; Fukuda, T.; Ashizuka, M.; Kohno, K.; Kuwano, M.; Sekiguchi, M. Binding capacity of human YB-1 protein for RNA containing 8-oxoguanine. Biochemistry 2002, 41 (42), 12739–12744. DOI: 10.1021/bi0201872 From NLM Medline.

(25) Hayakawa, H.; Sekiguchi, M. Human polynucleotide phosphorylase protein in response to oxidative stress. Biochemistry 2006, 45 (21), 6749–6755. DOI: 10.1021/bi052585l From NLM Medline.

(26) Hayakawa, H.; Fujikane, A.; Ito, R.; Matsumoto, M.; Nakayama, K. I.; Sekiguchi, M. Human proteins that specifically bind to 8-oxoguanine-containing RNA and their responses to oxidative stress. Biochem Biophys Res Commun 2010, 403 (2), 220–224. DOI: 10.1016/j.bbrc.2010.11.011 From NLM Medline.

(27) Ishii, T.; Hayakawa, H.; Igawa, T.; Sekiguchi, T.; Sekiguchi, M. Specific binding of PCBP1 to heavily oxidized RNA to induce cell death. Proc Natl Acad Sci U S A 2018, 115 (26), 6715–6720. DOI: 10.1073/pnas.1806912115 From NLM Medline.

(28) Seo, K. W.; Kleiner, R. E. YTHDF2 Recognition of N(1)-Methyladenosine (m(1)A)-Modified RNA Is Associated with Transcript Destabilization. ACS Chem Biol 2020, 15 (1), 132–139. DOI: 10.1021/acschembio.9b00655 From NLM Medline.

(29) Arguello, A. E.; DeLiberto, A. N.; Kleiner, R. E. RNA Chemical Proteomics Reveals the N(6)-Methyladenosine (m(6)A)-Regulated Protein-RNA Interactome. J Am Chem Soc 2017, 139 (48), 17249–17252. DOI: 10.1021/jacs.7b09213.

(30) Geuens, T.; Bouhy, D.; Timmerman, V. The hnRNP family: insights into their role in health and disease. Hum Genet 2016, 135 (8), 851–867. DOI: 10.1007/s00439-016-1683-5 From NLM Medline.

(31) Ishii, T.; Hayakawa, H.; Sekiguchi, T.; Adachi, N.; Sekiguchi, M. Role of Auf1 in elimination of oxidatively damaged messenger RNA in human cells. Free Radic Biol Med 2015, 79, 109–116. DOI: 10.1016/j.freeradbiomed.2014.11.018 From NLM Medline.

(32) Degrauwe, N.; Suva, M. L.; Janiszewska, M.; Riggi, N.; Stamenkovic, I. IMPs: an RNA-binding protein family that provides a link between stem cell maintenance in normal development and cancer. Genes Dev 2016, 30 (22), 2459–2474. DOI: 10.1101/gad.287540.116 From NLM Medline.

(33) Markus, M. A.; Morris, B. J. RBM4: a multifunctional RNA-binding protein. Int J Biochem Cell Biol 2009, 41 (4), 740–743. DOI: 10.1016/j.biocel.2008.05.027 From NLM Medline.

(34) Farina, K. L.; Huttelmaier, S.; Musunuru, K.; Darnell, R.; Singer, R. H. Two ZBP1 KH domains facilitate beta-actin mRNA localization, granule formation, and cytoskeletal attachment. J Cell Biol 2003, 160 (1), 77–87. DOI: 10.1083/jcb.200206003 From NLM Medline.

(35) Uniacke, J.; Holterman, C. E.; Lachance, G.; Franovic, A.; Jacob, M. D.; Fabian, M. R.; Payette, J.; Holcik, M.; Pause, A.; Lee, S. An oxygen-regulated switch in the protein synthesis machinery. Nature 2012, 486 (7401), 126–129. DOI: 10.1038/nature11055 From NLM Medline.

(36) Kazan, H.; Ray, D.; Chan, E. T.; Hughes, T. R.; Morris, Q. RNAcontext: a new method for learning the sequence and structure binding preferences of RNA-binding proteins. PLoS Comput Biol 2010, 6 (7), e1000832. DOI: 10.1371/journal.pcbi.1000832 From NLM Medline.

(37) Ray, D.; Kazan, H.; Chan, E. T.; Pena Castillo, L.; Chaudhry, S.; Talukder, S.; Blencowe, B. J.; Morris, Q.; Hughes, T. R. Rapid and systematic analysis of the RNA recognition specificities of RNA-binding proteins. Nat Biotechnol 2009, 27 (7), 667–670. DOI: 10.1038/nbt.1550 From NLM Medline.

(38) Foroushani, A. K.; Chim, B.; Wong, M.; Rastegar, A.; Smith, P. T.; Wang, S.; Barbian, K.; Martens, C.; Hafner, M.; Muljo, S. A. Posttranscriptional regulation of human endogenous retroviruses by RNA-binding motif protein 4, RBM4. Proc Natl Acad Sci U S A 2020, 117 (42), 26520–26530. DOI: 10.1073/pnas.2005237117 From NLM Medline.

(39) Ishii, T.; Sekiguchi, M. Two ways of escaping from oxidative RNA damage: Selective degradation and cell death. DNA Repair (Amst) 2019, 81, 102666. DOI: 10.1016/j.dnarep.2019.102666 From NLM Medline.

(40) Barreau, C.; Paillard, L.; Osborne, H. B. AU-rich elements and associated factors: are there unifying principles? Nucleic Acids Res 2005, 33 (22), 7138–7150. DOI: 10.1093/nar/gki1012 From NLM Medline.

(41) Jonson, L.; Vikesaa, J.; Krogh, A.; Nielsen, L. K.; Hansen, T.; Borup, R.; Johnsen, A. H.; Christiansen, J.; Nielsen, F. C. Molecular composition of IMP1 ribonucleoprotein granules. Mol Cell Proteomics 2007, 6 (5), 798–811. DOI: 10.1074/mcp.M600346-MCP200 From NLM Medline.

(42) Moraes, K. C.; Quaresma, A. J.; Maehnss, K.; Kobarg, J. Identification and characterization of proteins that selectively interact with isoforms of the mRNA binding protein AUF1 (hnRNP D). Biol Chem 2003, 384 (1), 25–37. DOI: 10.1515/BC.2003.004 From NLM Medline.

(43) Wang, Y.; Chen, D.; Qian, H.; Tsai, Y. S.; Shao, S.; Liu, Q.; Dominguez, D.; Wang, Z. The splicing factor RBM4 controls apoptosis, proliferation, and migration to suppress tumor progression. Cancer Cell 2014, 26 (3), 374–389. DOI: 10.1016/j.ccr.2014.07.010 From NLM Medline.

(44) Kar, A.; Havlioglu, N.; Tarn, W. Y.; Wu, J. Y. RBM4 interacts with an intronic element and stimulates tau exon 10 inclusion. J Biol Chem 2006, 281 (34), 24479–24488. DOI: 10.1074/jbc.M603971200 From NLM Medline.

(45) Lin, J. C.; Tarn, W. Y. Exon selection in alpha-tropomyosin mRNA is regulated by the antagonistic action of RBM4 and PTB. Mol Cell Biol 2005, 25 (22), 10111–10121. DOI: 10.1128/MCB.25.22.10111-10121.2005 From NLM Medline.

(46) Lin, J. C.; Hsu, M.; Tarn, W. Y. Cell stress modulates the function of splicing regulatory protein RBM4 in translation control. Proc Natl Acad Sci U S A 2007, 104 (7), 2235–2240. DOI: 10.1073/pnas.0611015104 From NLM Medline.

