## Supplementary Information for "Chemoproteomic profiling of 8-oxoguanosine-sensitive RNA-protein interactions"

Contents

| | |
|---|---|
| Materials and Methods | 2 |
| Supplementary Tables | 7 |
| Supplementary Figures | 8 |

### Materials and Methods

*Plasmids*

Plasmids encoding cDNA were obtained from Addgene: IGF2BP1 (#21659), IGF2BP2 (#91890), HNRNPD (#38066); purchased from Genscript: RBM4 (#OHu17642D), RBM4B (#OHu04653D); or purchased from Origene: HNRNPDL (#SC107613). For mammalian transfection, sequences were cloned into a modified pcDNA5/FRT/TO with an N-terminal 3xFLAG tag. For protein expression and purification in *E. coli*, constructs were cloned into pGEX-6P-1, pET21a or a modified pET28a vector.

*Cell culture*

HeLa and HEK293T cells were cultured at 37°C in a humidified atmosphere with 5% $CO_2$ in DMEM (Life Technologies) supplemented with 10% FBS (Atlanta), 1× penicillin-streptomycin and 2 mM L-glutamine (Life Technologies). For transfection experiments, 3 million HEK293T cells were plated in a 10 cm dish 16 hours prior to $CaPO_4$ transfection. 10 µg of plasmid DNA was used per 10 cm dish. Cells were harvested 24 hours later.

*Oligonucleotide synthesis*

Diazirine-containing G or 8OG oligonucleotide probes were synthesized as previously described[1]. In brief, 5-aminoallyluridine-modified RNA oligonucleotides were purchased from Dharmacon and reacted with diazirine NHS-ester. Labeled product was purified by reverse-phase HPLC on a Zorbax Eclipse XDB-C18 semipreparative column using an Agilent 1260 Infinity system. Characterization was performed by high-resolution mass spectrometry (HRMS) on an Agilent 6220 Accurate-Mass Time-of-Flight LC/MS (ESI-TOF) in negative mode. 3'-fluorescein-modified oligonucleotides were synthesized by solid-phase oligonucleotide synthesis on an Applied Biosystems (ABI) 394 automated synthesizer using 3'-(6-fluorescein) CPG (Glen

Research) and 8-oxoguanosine phosphoramidite (ANP-9420, ChemGenes) following the manufacturer's recommendations. Oligonucleotides were purified by HPLC and confirmed by HRMS.

*Photocrosslinking reactions with cellular lysates*

For proteomics experiment, HeLa S3 cell lysates were photocrosslinked with RNA probes and crosslinked RNA-protein complexes were isolated by streptavidin enrichment, as previously described[1]. In brief, HeLa S3 cells were harvested and lysed by cryomilling. The resulting cell powder was first extracted with low-salt extraction buffer (20 mM Tris HCl pH 7.5, 10 mM NaCl, 2 mM $MgCl_2$, 0.5% Triton X-100, 10% glycerol, protease and phosphatase inhibitor tablet) and then with high-salt extraction buffer (50 mM Tris HCl pH 7.5, 420 mM NaCl, 2 mM $MgCl_2$, 0.5% Triton X-100, 10% glycerol, protease and phosphatase inhibitor tablet). High-salt and low-salt extracts were pooled and diluted to 2 mg/mL before proceeding to photocrosslinking. Photocrosslinking reactions were performed in a 60 mm dish using 2 mL of extract and 1 μM RNA oligo probe. Reactions were incubated on ice for 20 minutes and crosslinked on ice for 15 minutes with 365 nm UV (Spectroline ML-3500S). Each reaction was incubated with 60 μl of 50% streptavidin agarose bead slurry (Pierce #20357) for 3 hours at 4°C with end-to-end rotation. Beads were washed three times with 1% SDS in 1X TBS, three times with 6M urea in 1X TBS, and three times with 1X TBS. RNA-bound proteins were eluted with 50 μL of RNase elution buffer (10 mM Tris HCl pH 7.5, 40 mM NaCl, 1 mM $MgCl_2$, 25 unit/mL RNase A, and 2000 unit/mL RNase T1) at 37°C for 30 minutes with periodic mixing. For western blotting, wild type or transfected HEK293T cells were lysed in NP-40 lysis buffer (50 mM Tris-HCl pH 7.5, 150 mM NaCl, 0.5% Nonidet P-40, 5 mM $MgCl_2$, 1 mM PMSF, and protease inhibitor tablet). Clarified lysates were incubated with RNA probes on ice for 20 minutes and photocrosslinked on ice for 10 minutes with 365 nm UV. Cross-linked RNA-protein complexes were incubated with streptavidin agarose beads, washed three times with 1% SDS in 1X TBS, and eluted with SDS sample buffer at 95°C for 5 minutes. Input and elution samples were loaded on an SDS-PAGE gel and analyzed by western blotting.

*Mass spectrometry proteomics*

Proteomics analysis was performed as previously described[1]. Quantification of protein abundance was performed by spectral counting, and one spectral count was added to all values to enable calculation of fold change. Proteins were only selected for quantification if they were present in at least three out of six samples, and data were normalized using Scaffold. A student's t-test was used to determine statistical significance. Protein fold-change was determined by calculating the geometrical mean of the protein fold-changes across three biological replicates.

*Western blotting*

For western blot analysis, the following antibodies were used: anti-FLAG M2 (Sigma Aldrich #F3165 used at a 1:1000 dilution), anti-HNRNPD (kind gift from Professor Steitz, New Haven, CT, used at a 1:2000 dilution), anti-IGF2BP1 (kind gift from Professor Hüttelmaier, Halle, Germany, used at a 1:1000 dilution). IRDye-conjugated secondary antibodies raised in donkey (anti-mouse 680LT and anti-rabbit 800CW) and IRDye-conjugated streptavidin (streptavidin 800CW) were purchased from LI-COR Biosciences and used according to manufacturer's instructions.

*Protein expression and purification*

Proteins were expressed in *Escherichia coli* strain BL21 (Rosetta). IGF2BP1-KH (195-577), IGF2BP1-RRM (1-194) and full-length IGF2BP2 were purified as previously described[2]. In brief, the GST-tagged proteins were affinity purified using Glutathione Sepharose 4B resin (GE Healthcare). IGF2BP1-KH and IGF2BP1-RRM were eluted from the beads with glutathione, while full-length IGF2BP2 was eluted by GST-PreScission protease and further purified by linear salt elution from Heparin column (GE Healthcare). GST-tagged RBM4-RRM (1-176) was expressed overnight at 18°C with 0.2 mM IPTG and affinity purified using a 1 ml GSTrap 4B column (GE Healthcare). RBM4-RRM was eluted from the beads with glutathione.

*EMSA*

Fluorescein-labeled RNA oligonucleotides (100 nM) were titrated with 3-fold protein dilutions ranging from 9 μM to 40 nM in reaction buffer containing 10 mM Tris-HCl pH 7.5, 50 mM NaCl, 1 mM $MgCl_2$, 0.5 mM EDTA, 1 mM DTT and 4% glycerol. The reaction mix was incubated at room temperature for 30 min, then glycerol was added to reach a final concentration of 20% and the mix was further incubated on ice for 15 minutes. RNA-protein complexes were resolved on a 5% TGE native gel and imaged using an ImageQuant LAS 4000 fluorescence imager. The proportion of non-complexed RNA was measured by densitometry by drawing an identically sized box around each band and measuring the raw integrated density minus background. These values were then fit to a four-parameter dose-response curve (GraphPad Prism). Intensity values were normalized and converted to percent bound.

*Fluorescence polarization*

Fluorescein-containing RNA oligonucleotides (20nM) were titrated with 3-fold protein dilutions ranging from 30 μM to 1 nM in reaction buffer containing 50 mM Tris-HCl pH 7.5, 150 mM NaCl, 5 mM $MgCl_2$, 0.5% NP-40, mM EDTA, 1 mM DTT. The reaction mix was incubated at room temperature for 30 minutes and fluorescence polarization was measured using an EnVision multimode plate reader. Fluorescence polarization values were then fit to a four-parameter dose-response curve (GraphPad Prism).

*Photocrosslinking with purified proteins*

Purified proteins (100-200 nM) were titrated with 3-fold RNA probe dilutions ranging from 3 μM to 4 nM in reaction buffer containing 20 mM Tris pH 7.5, 150 mM NaCl. The reaction mix was incubated on ice for 20 minutes and photo-cross-linked on ice for 10 minutes with 365 nm UV. Crosslinked RNA-protein complexes were resolved by SDS-PAGE and analyzed by streptavidin western blotting. Complex formation was measured by densitometry as described for EMSA. Values were then fit to a four-parameter dose-response curve (GraphPad Prism). Intensity values were normalized and converted to crosslinking efficiency (%), where 100% crosslinking efficiency corresponds to the highest signal on each blot.

**Supplementary Table 1.** RNA oligonucleotides used in this work.

| Oligo | RNA Sequence | Calculated $[M\text{-}3H]^{3-}$ | Measured $[M\text{-}3H]^{3-}$ |
|---|---|---|---|
| **1** | 5’-AC-8OG-UG-5DzU-GUAC-biotin-3’ | 1302.24 | 1302.24 |
| **2** | 5’-ACGUG-5DzU-GUAC-biotin-3’ | 1296.91 | 1296.92 |
| **3** | 5’-GA-8OG-CU-5DzU-GUAC-biotin-3’ | 1302.24 | 1302.24 |
| **4** | 5’-GAGCU-5DzU-GUAC-biotin-3’ | 1296.91 | 1296.91 |
| **5** | 5’-AC-8OG-UGUGUAC-fluorescein-3’ | 1246.52 | 1246.51 |
| **6** | 5’-ACGUGUGUAC-fluorescein-3’ | 1241.19 | 1241.19 |
| **7** | 5’-GUAAC-8OG-GUAC-fluorescein-3’ | 1254.19 | 1254.19 |
| **8** | 5’-GUAACGGUAC-fluorescein-3’ | 1248.86 | 1248.86 |
| **9** | 5’-CGCG-8OG-CGGUA-fluorescein-3’ | 1264.53 | 1264.53 |
| **10** | 5’-CGCGGCGGUA-fluorescein-3’ | 1259.20 | 1259.21 |

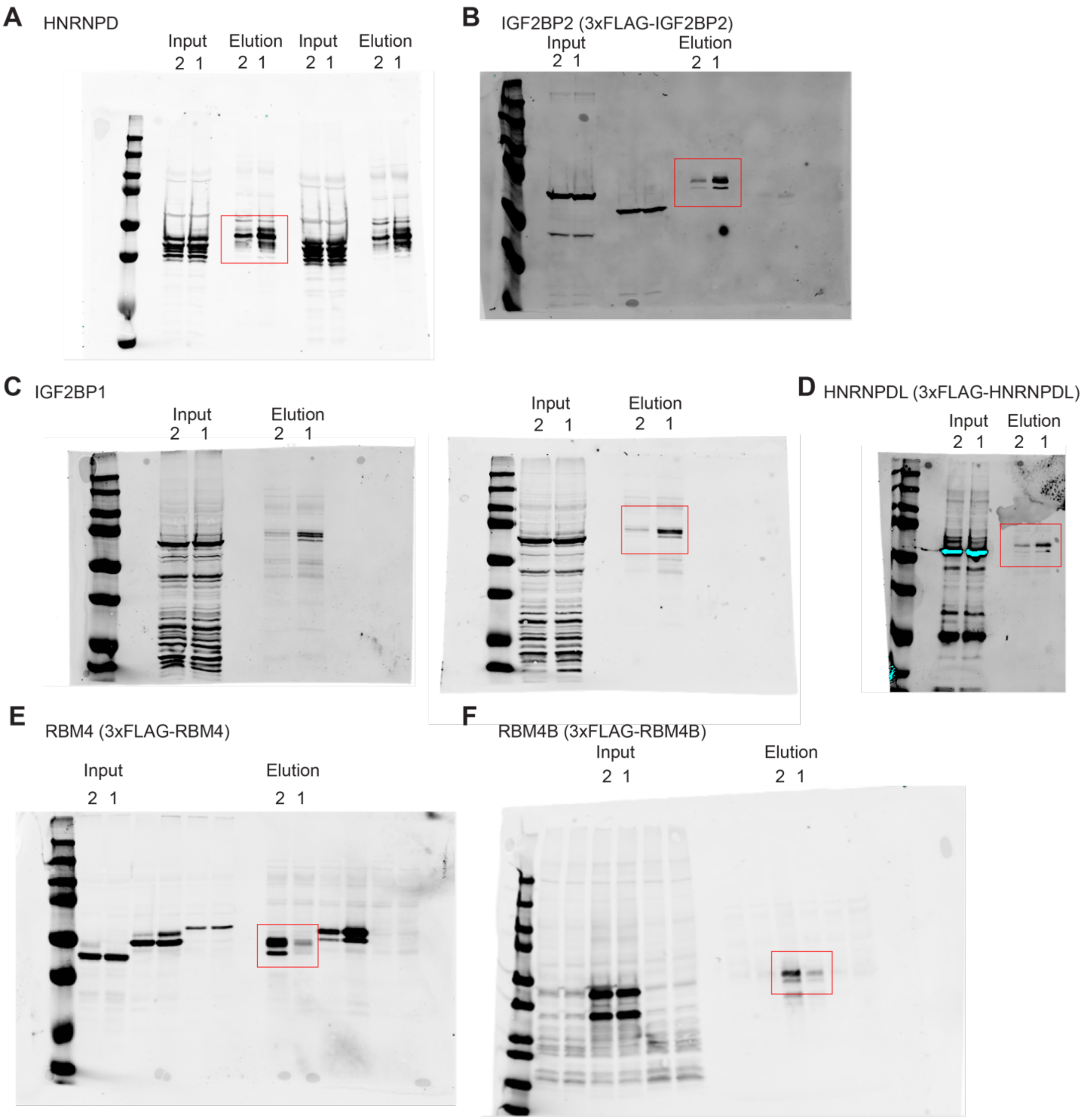


**Supplementary Figure 1.** Western blot validation of proteomics results. Full western blots for Figure 1C. Cell lysates from either HEK293T or HEK293T cells transfected with 3xFLAG-tagged hnRNPDL, IGF2BP2 or RBM4/4B were photocrosslinked with 1 µM oligo probe **1** (8OG) or **2** (G), streptavidin enriched, and analyzed using antibodies against HNRNPD **(A)**, IGF2BP1 **(C)** or anti-FLAG antibodies **(B, D, E, F)**.

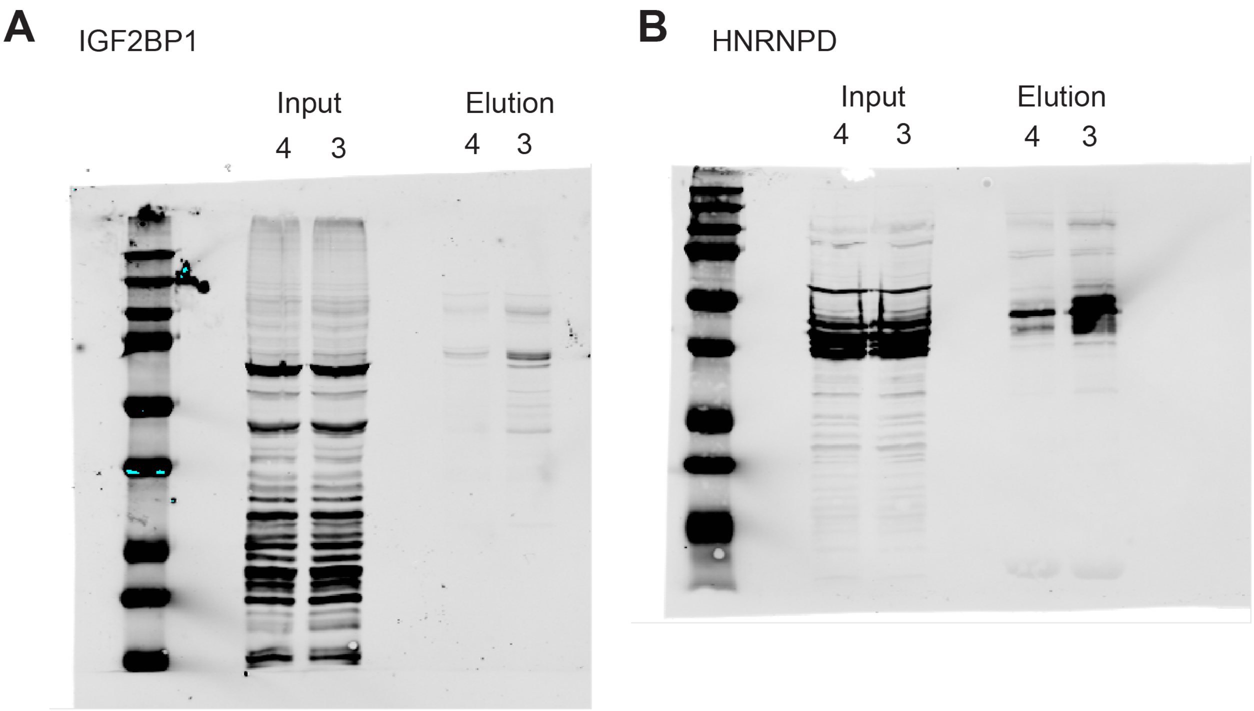


**Supplementary Figure 2.** Pulldown of IGF2BP1 and HNRNPD with scrambled sequence oligo probes **3** (8oG) or **4** (G). Cell lysates were photocrosslinked as in Figure 1C and analyzed using antibodies against IGF2BP1 **(A)** or HNRNPD **(B)**.

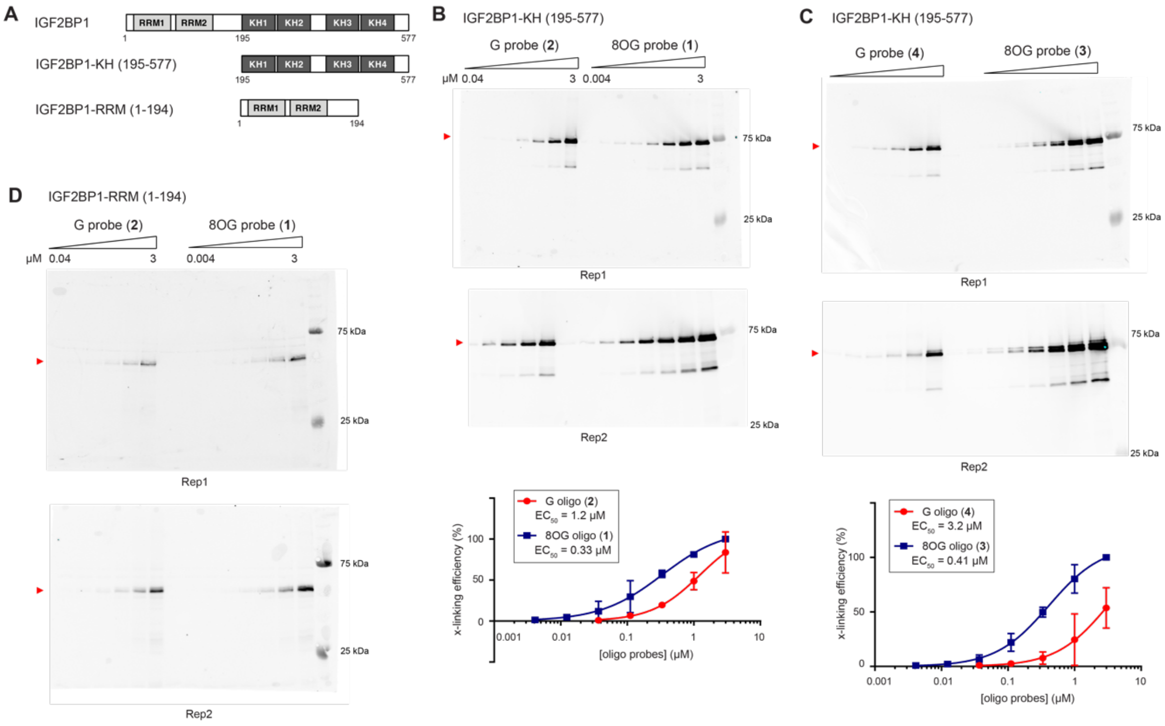


**Supplementary Figure 3.** Photocrosslinking of IGF2BP1 KH and RRM domains with G or 8OG oligo probes. **(A)** IGF2BP1 recombinant protein constructs expressed in *E. coli.* **(B)** 100 nM recombinant IGF2BP1-KH (195-577) was photocrosslinked with oligo probes **1** or **2** at concentrations ranging from 1 nM to 3 μM. Crosslinked species were detected by streptavidin western blotting. **(C)** 100 nM recombinant IGF2BP1-KH (195-577) was photocrosslinked with oligo probes **3** or **4** as in **(B)**. **(D)** IGF2BP1-RRM (1-194) was photocrosslinked with oligo probes **1** or **2** as in **(B)**.

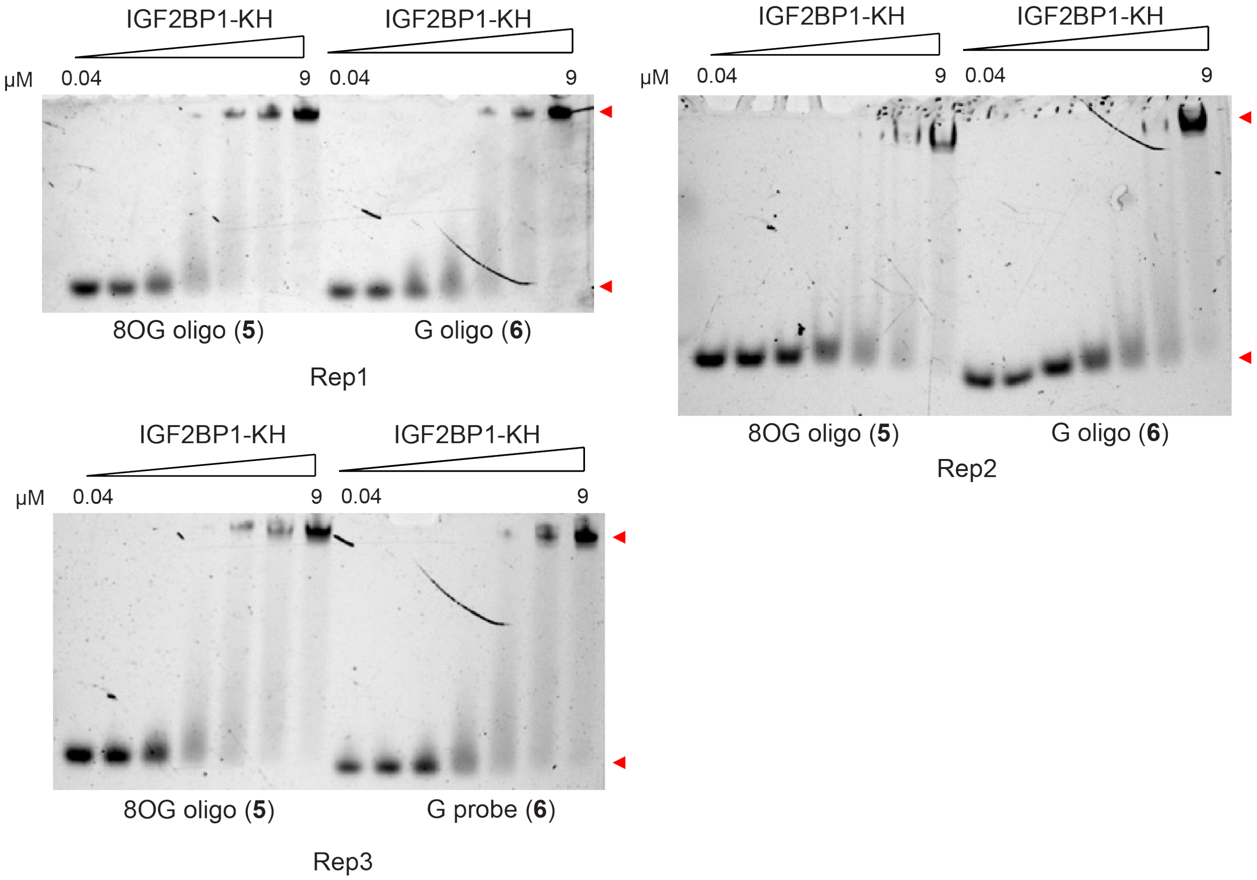


**Supplementary Figure 4.** Characterization of the interaction between IGF2BP1-KH (195-577) and fluorescein-labeled oligos **5** or **6** by EMSA. Full data for Figure 2A in the main text. 100 nM oligo was incubated with IG2BP1-KH at concentrations ranging from 40 nM to 9 µM in low salt reaction buffer and resolved on a Tris-glycine-EDTA (TGE) native gel.

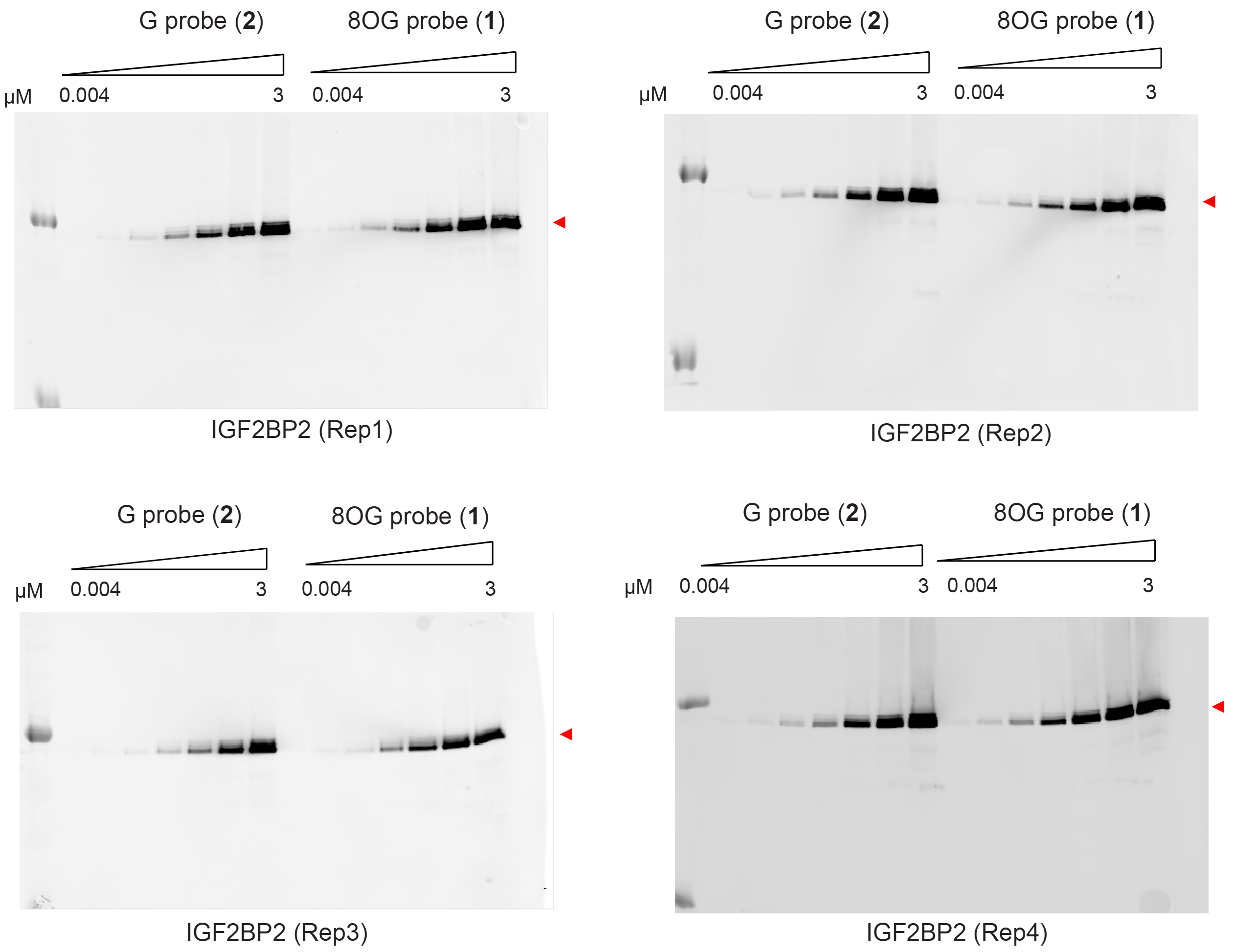


**Supplementary Figure 5.** Photocrosslinking between recombinant IGF2BP2 and G or 8OG probes **1** and **2**. Full blot data for Figure 2C in the main text. Experiment was conducted as described in Supplementary Figure 3.

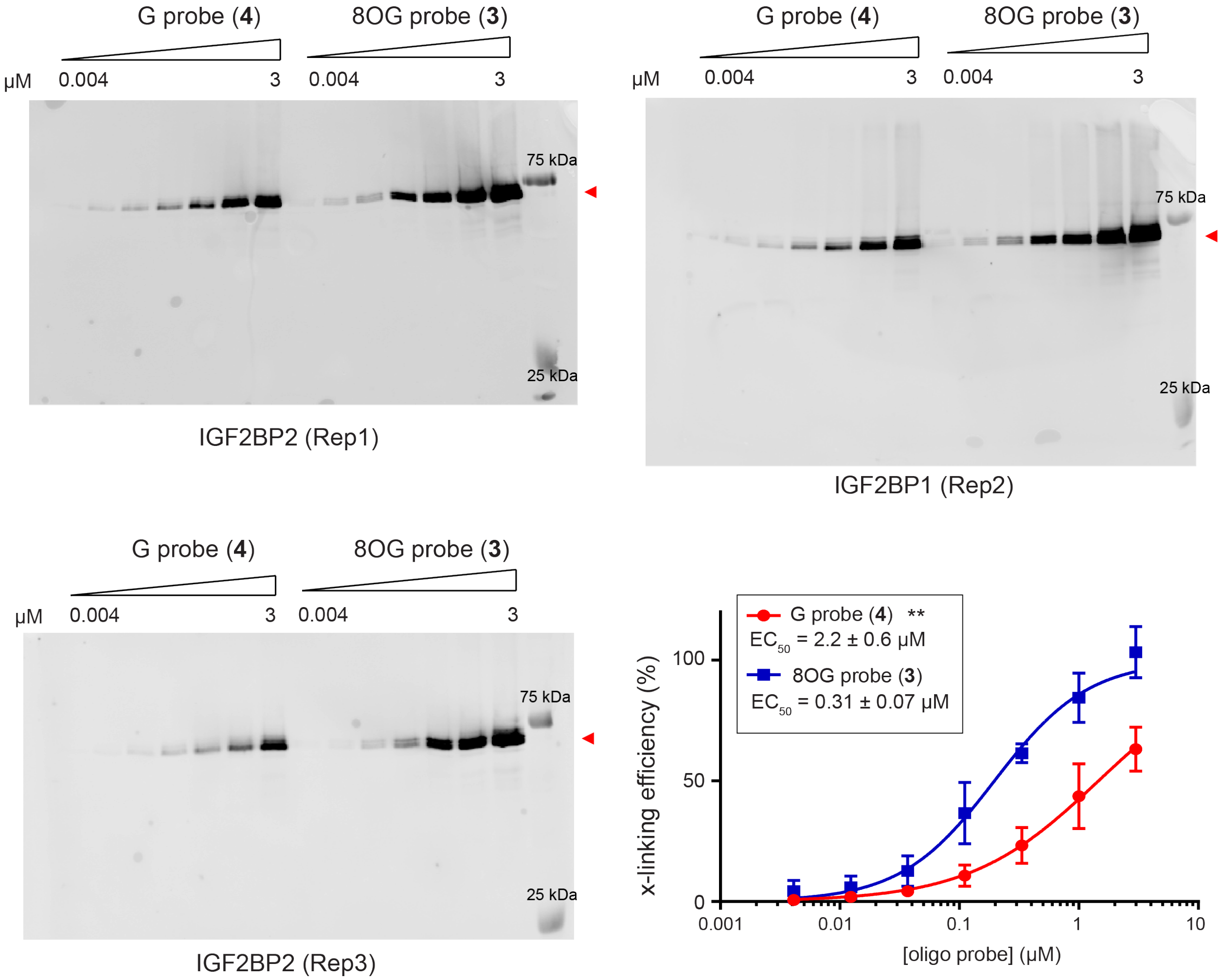


**Supplementary Figure 6.** Photocrosslinking between recombinant IGF2BP2 and G or 8OG probes **3** and **4**. Experiment was conducted as described in Supplementary Figure 3.

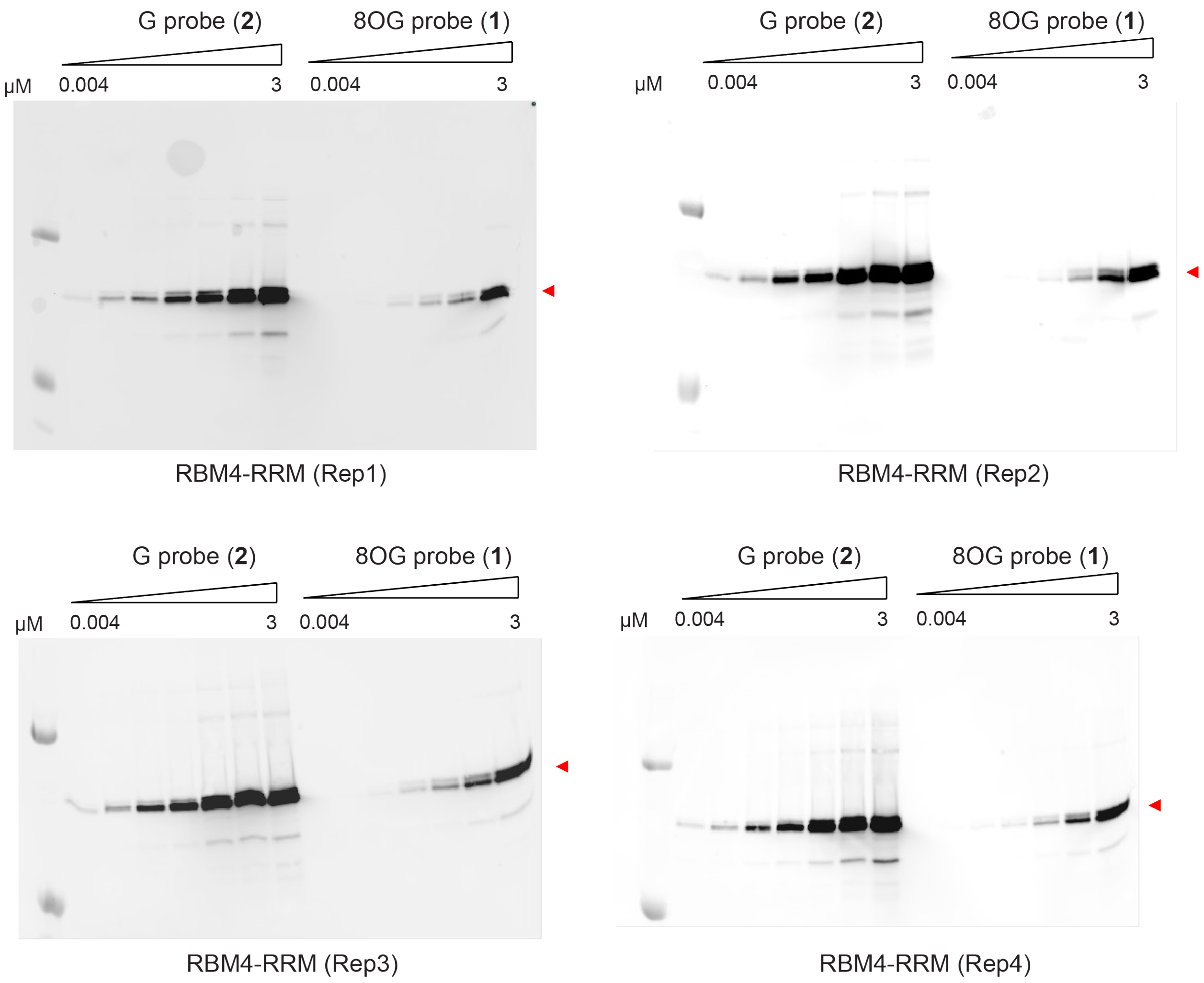


**Supplementary Figure 7.** Photocrosslinking between recombinant RBM4-RRM (1-176) and G or 8OG probes **1** and **2.** Full blot data for Figure 3A in the main text. Experiment was conducted as described in Supplementary Figure 3 except that 200 nM of protein was used.

### References

(1) Arguello, A. E.; DeLiberto, A. N.; Kleiner, R. E. RNA Chemical Proteomics Reveals the N(6)-Methyladenosine (m(6)A)-Regulated Protein-RNA Interactome. *J Am Chem Soc* **2017**, *139* (48), 17249-17252. DOI: 10.1021/jacs.7b09213.

(2) Wachter, K.; Kohn, M.; Stohr, N.; Huttelmaier, S. Subcellular localization and RNP formation of IGF2BPs (IGF2 mRNA-binding proteins) is modulated by distinct RNA-binding domains. *Biol Chem* **2013**, *394* (8), 1077-1090. DOI: 10.1515/hsz-2013-0111 From NLM Medline.
